# Natural selection reverses the exaggeration of a male sexually selected trait, which increases female fitness

**DOI:** 10.1101/2020.10.15.340562

**Authors:** Kensuke Okada, Masako Katsuki, Manmohan D. Sharma, Katsuya Kiyose, Tomokazu Seko, Yasukazu Okada, Alastair J. Wilson, David J. Hosken

**Affiliations:** Graduate School of Environmental and Life Science, 111 Tsushima-naka, Okayama City, Okayama 700-8530, Japan; Department of Agricultural and Environmental Biology, The University of Tokyo, Yayoi 1-1-1, Bunkyo-ku, Tokyo 113-8657, Japan; Centre for Ecology & Conservation, University of Exeter, Cornwall, UK; National Agriculture and Food Research Organization, Central Region Agricultural Research Center, Kannondai 2-1-18, Tsukuba, Ibaraki 305-8666, Japan; Department of Biological Sciences, School of Science, Tokyo Metropolitan University, Minamiohsawa 1-1, Hachiohji, Tokyo 192-0397, Japan

**Keywords:** sexual selection, predation, sexual conflict, genetic correlation, fitness

## Abstract

Theory shows how sexual selection can exaggerate male traits beyond naturally selected optima and also how natural selection can ultimately halt trait elaboration. Empirical evidence supports this theory, but to date, there have been no experimental evolution studies directly testing this logic, and little examination of possible associated effects on female fitness. Here we used experimental evolution of replicate populations of broad-horned flour-beetles to test for evolutionary effects of sex-specific predation on an exaggerated sexually selected male trait, while also testing for effects on female lifetime reproductive success. We found that populations subjected to male-specific predation evolved smaller sexually selected traits and this indirectly increased female fitness, seemingly through intersexual genetic correlations we documented. Predation solely on females had no effects. Our findings support fundamental theory, but also reveal novel outcomes when natural selection targets sex-limited sexually selected characters.

## Introduction

Sexual selection typically acts more strongly on males and is responsible for the evolution of a vast array of exaggerated characters that enhance male sexual fitness components (Andersson 1994; Shuster & Wade 2003; Andersson & Simmons 2006; Hosken & House 2011). Lande’s (1981) and Kirkpatrick’s (1982) models of sexual selection via the Fisher (1930) process – the null models of intersexual selection (Prum 2010) – clearly shows how this can occur. They also demonstrates how natural selection can oppose sexual selection as trait values move beyond their naturally selected optima (reviewed in Arnold 1983). While theory is clear on the joint effects of natural and sexual selection on sexual trait evolution, explicit experimental tests of theoretical predictions are required to fully understand sexual trait evolution (Pocklington & Dill 1995).

One potentially important source of natural selection that could affect the evolution of sexually selected traits is predation (Andersson 1994), and many studies have shown predation can seemingly oppose the exaggeration of male sexual characters. For example, sexual signals are conspicuous to potential mates but may also attract predators and parasitoids (Zuk & Kolluru 1998). This is well documented in orthopterans and frogs (e.g. Cade 1975; Ryan et al. 1982; Sakaluk & Belwood 1984; Ryan 1985; Hosken et al. 1994; Rotenbury et al. 1996; Gray & Cade 1999), and this form of natural selection is probably responsible for the loss of cricket sexual signals on two Hawaiian islands (Zuk et al. 2006; Pascoal et al. 2014). More generally, predation appears reverse the evolution of extreme sexually selected phenotypes (reviewed in Kotiaho 2001) and males frequently reduce their sexual signaling in response to predation risk, which can result in decreased mating success when risk is high (Lima and Dill 1990; Sakaluk 1990; Candolin 1998; Kotiaho et al. 1998). Nonetheless, while there is ample evidence that predation selects against sexual trait enhancement, there is limited direct experimental verification that this generates evolutionary responses in these characters (but see e.g. Zuk et al. 1993; Millar et al. 2006).

Female reproductive success can also be impacted by predation (Reznick et al. 1990; Magnhagen 1991). For example, egg-carrying females can be slower (Cooper et al. 1990; Berglund and Rosenqvist 1986) and suffer higher predation rates (Lee et al. 1996; Magnhagen 1991). Resultant anti-predator behaviours may reduce foraging efficiency and reproductive activity, and thus be costly to females (reviewed in Lima and Dill 1990). Costs can be generated by delayed development, slower growth or postponed reproduction (Barry 1994; Clobert et al. 2000; Koskela and Ylönen 1995; Lind & Cresswell 2005). Nonetheless, while there is ample evidence that predation selects on both sexes, selection on females and males is frequently investigated independently, and again the evolutionary effects of any sex-specific selection are frequently inferred. That is, controlled experimental tests of evolutionary responses to this selection are usually not undertaken. Unfortunately, without exploring how predation affects both sexes, we are unlikely to fully understand how predation affects sexual trait evolution (Pocklington & Dill 1995). This is especially true when intersexual genetic correlations link sexually selected male characters with female fitness because selection on one sex can affect the other via these correlations.

Here we tested for effects of predation on the evolution of a male sexually selected character and female lifetime reproductive success (LRS) in the broad-horned flour beetle *Gnatocerus cornutus*. Male beetles develop enormously enlarged mandibles that are used in male-male fights, and males with larger mandibles have higher fighting and mating success (Okada & Miyatake 2009; Harano et al. 2010). Females lack this exaggerated character completely (Okada et al. 2006). We have previously shown that males with larger mandibles sire daughters with lower fecundity, and that selecting for increased (or decreased) mandible size results in decreased (or increased) female fitness (LRS) (Harano et al. 2010). This apparently occurs because of the beetle’s genetic architecture, which means evolving larger mandibles results in the correlated evolution of masculinized females (even though females never develop mandibles). Basically, the enlarged male mandible requires a masculinised head and prothorax to function optimally and this means males with larger mandibles have smaller abdomens (Harano et al. 2010; Okada et al. 2012). Thus although females never develop mandibles, selecting on male mandibles ultimately affects female abdomen size, which likely determines the number of eggs a female carries (Harano et al. 2010, and also see Honĕk 1993). These previous *G. cornutus* studies clearly document intralocus sexual conflict over beetle morphology (e.g. Harano et al. 2010; Katsuki et al. 2012b) and point to a negative intersexual correlation between (male) mandible size and female fitness, although this link has not been directly established and requires confirmation (c.f. Pennell et al. 2016). With this in mind we investigated the beetle’s intersexual genetic architecture for key focal traits using an animal model (Wilson et al. 2010). We also established replicate (3/treatment) experimental evolution populations subjected to either male or female predation, along with control populations (n = 3, for 9 total populations). After 8 generations of experimental evolution, we assessed female fitness (LRS) and measured a range of morphological characters, including mandible size. Morphology was also measured during experimental evolution. We found strong effects of male-specific predation on morphology and female fitness, while predation on females alone had no effects.

## Results

The genetic parameters estimated from the animal model analyses confirmed what previous experimental evolution studies had inferred (e.g. Okada & Miyatake 2009; Harano et al. 2010). Likelihood ratio comparisons of univariate models confirmed the presence of additive genetic variance in all four traits (supplemental materials) as expected. Note that the decision to treat the two mass measures as the same trait across the sexes was justified by an absence of significant genotype-by-sex interaction in univariate models. Cross-sex genetic correlations were estimated as c.+1 for both traits, but we do note there is a qualitative pattern of higher additive variance for body mass in females (see supplemental materials for these analyses and some further comment).

Multivariate models further confirmed the presence of additive genetic variance (LRT comparison of null model to one with a diagonal genetic matrix; χ^2^_4_=74.96, P<0.0001), as well as significant among-trait genetic correlation structure (LRT comparison of model with a diagonal genetic matrix to full model with all genetic correlations included; χ^2^_6_=39.02, P<0.0001). Parameter estimates from the full multivariate model show substantial genetic variation in all traits measured and reasonably high trait heritability (Table 1). All traits were positively genetic correlated except male mandible size and female fitness (lifetime reproductive success) and male mandible size and abdominal mass, which were both strongly negatively correlation. Individual genetic correlations were nominally significant at α=0.05 (based on |r_G_| > 1.96SE) except between body and abdomen mass and between body mass and female LRS.

**Table 1.** Estimates of the genetic variance-covariance structure among traits in the breeding design. Estimates are shown standardized to narrow sense heritability (h^2^; shaded diagonal) and genetic correlations (r_G_; above diagonal). Values were estimated using a four trait animal model with body and abdomen mass treated as the same trait in both sexes (but with sex as a fixed effect). Standard errors are denoted in parentheses and bold font denotes estimates that are nominally significant at P<0.05 assuming approximate 95% CI are provided by the estimate ± 1.96 SE.

| Traits | Body mass | Abdominal mass | Male mandible size | Female LRS |
| --- | --- | --- | --- | --- |
| Body mass | **0.346** (0.085) | 0.296 (0.164) | **0.574** (0.162) | 0.422 (0.242) |
| Abdominal mass |  | **0.513** (0.096) | **-0.415** (0.171) | **0.596** (0.199) |
| Male mandible size |  |  | **0.380** (0.111) | **-0.598** (0.250) |
| Female LRS |  |  |  | **0.214** (0.106) |

The experimental treatments we imposed on the replicated beetles populations generated treatment specific evolutionary responses in some traits but not others (Figure 1). When we compared trait values at the end of the experimental evolution period we found that predation significantly affected male mandible size (Figure 2a. *F*_2,6_ = 31.07; *P* < 0.001). *Post-hoc t*-tests [with sequential Bonferroni adjustment] revealed that when males were exposed to predators they evolved the smallest mandibles (all *P* < 0.01), while the control and female treatments did not differ in mandible size (*P* = 0.36). Similarly, predation effected male abdomen size (*F*_2,6_ = 31.04; *P* < 0.001), and *post-hoc t-*test revealed that males exposed to predators evolved the largest abdomens (all *P* < 0.01) while control and female predator-exposure treatments did not differ (*P* = 1.0) (Figure 2b). Total male body size was unaffected by our treatments (Figure 2c: *F*_2,6_ = 1.17; *P* = 0.373).

**Figure 1.**
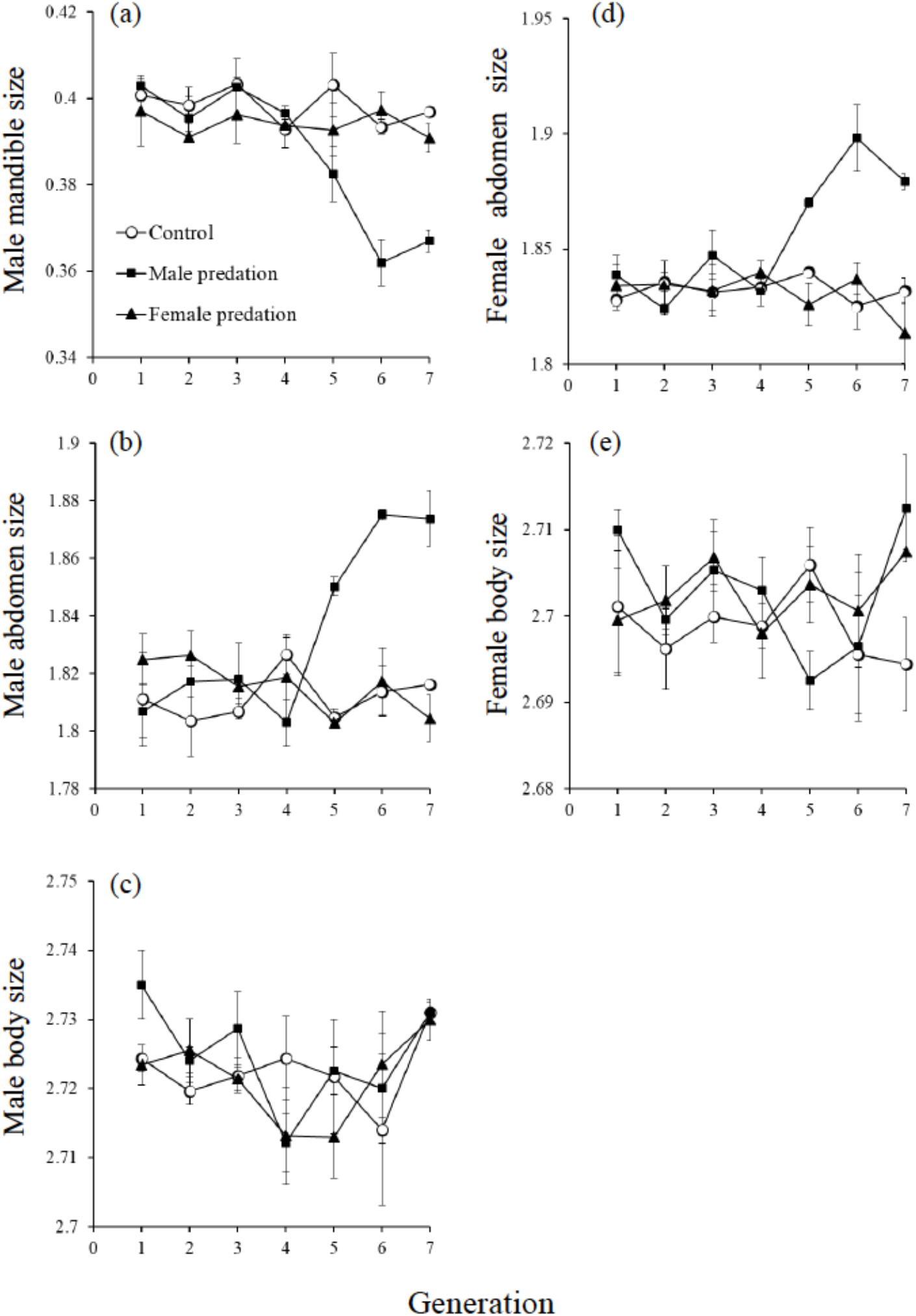
Responses to selection on predation in male mandible size (mm) (a), male abdomen size (mg) (b), male body size (mg) (c), female abdomen size (mg) (d) and female body size (mg) (e) (mean ± SE). White circles are the control populations that were not subjected to selection by predation. Black squares and triangles, are the populations with male and female exposure to predators, respectively. Note we did not measure female fitness (lifetime reproductive success: LRS) at every generation as it was not logistically possible.

**Figure 2.**
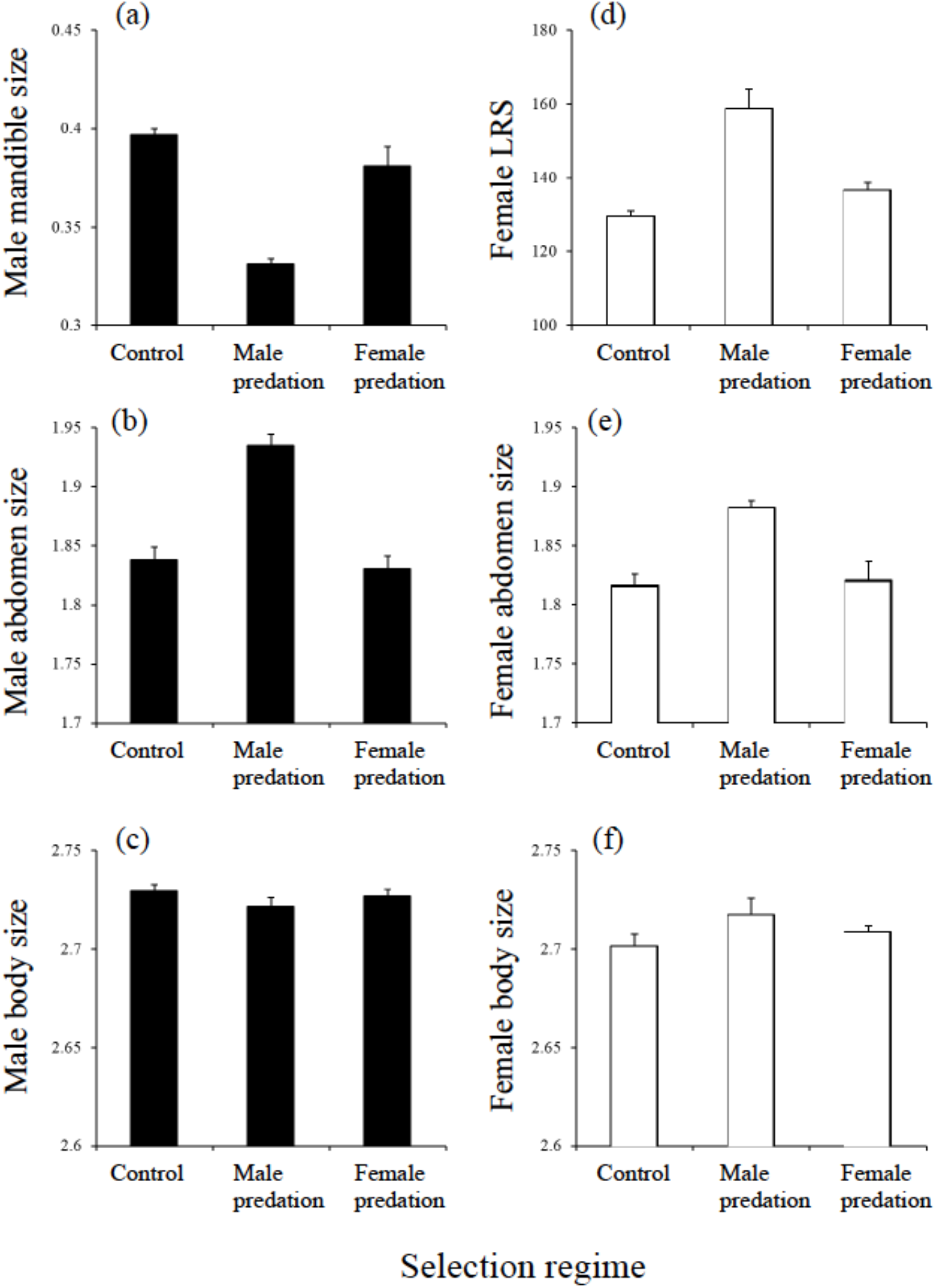
Trait values after 8 generation of experimental evolution. Replicate populations (3/treatment) were exposed to either male-only predation (middle columns), female-only predation (right-hand columns) or no predation (controls: left-hand columns), and effects on a range of traits was assessed. Traits were: Male mandible size (mm) (a), Male abdomen size (mg) (b), Male body size (mg) (c), Female lifetime reproductive success (LRS: offspring number) (d), Female abdomen size (mg) (e) and Female body size (mg) (f) (shown are means (population as replicate) ± SE).

Predation also effected female LRS (Figure 2d: *F*_2,6_ = 21.29; *P* = 0.002). *Post-hoc t*-tests showed that this was because in the male predation treatment female fitness was higher (all *P* < 0.01), while the other treatments did not differ (control = female predation: *P* = 0.54). Thus exposing males to predators resulted in the evolution of smaller male mandibles and higher female fitness. As with males, female abdomen size was impacted by our treatments (*F*_2,6_ = 10.75; *P* = 0.010) and again the male predator exposure group evolved the largest female abdomens (all *P* ≤ 0.01: control = female predation *P* = 1.0). Finally, female body size was unaffected by our experimental regimes (*F*_2,6_ = 1.65; *P* = 0.269).

To summarise the main findings: male-specific predation results in an altered, more feminized male phenotype, that included reductions in the size of a male-limited sexually selected trait. Additionally, because of the beetle’s genetic architecture, the “new” demasculinized male phenotype was transmitted through to the female phenotype and resulted in higher fitness females (Figure 3).

**Figure 3.**
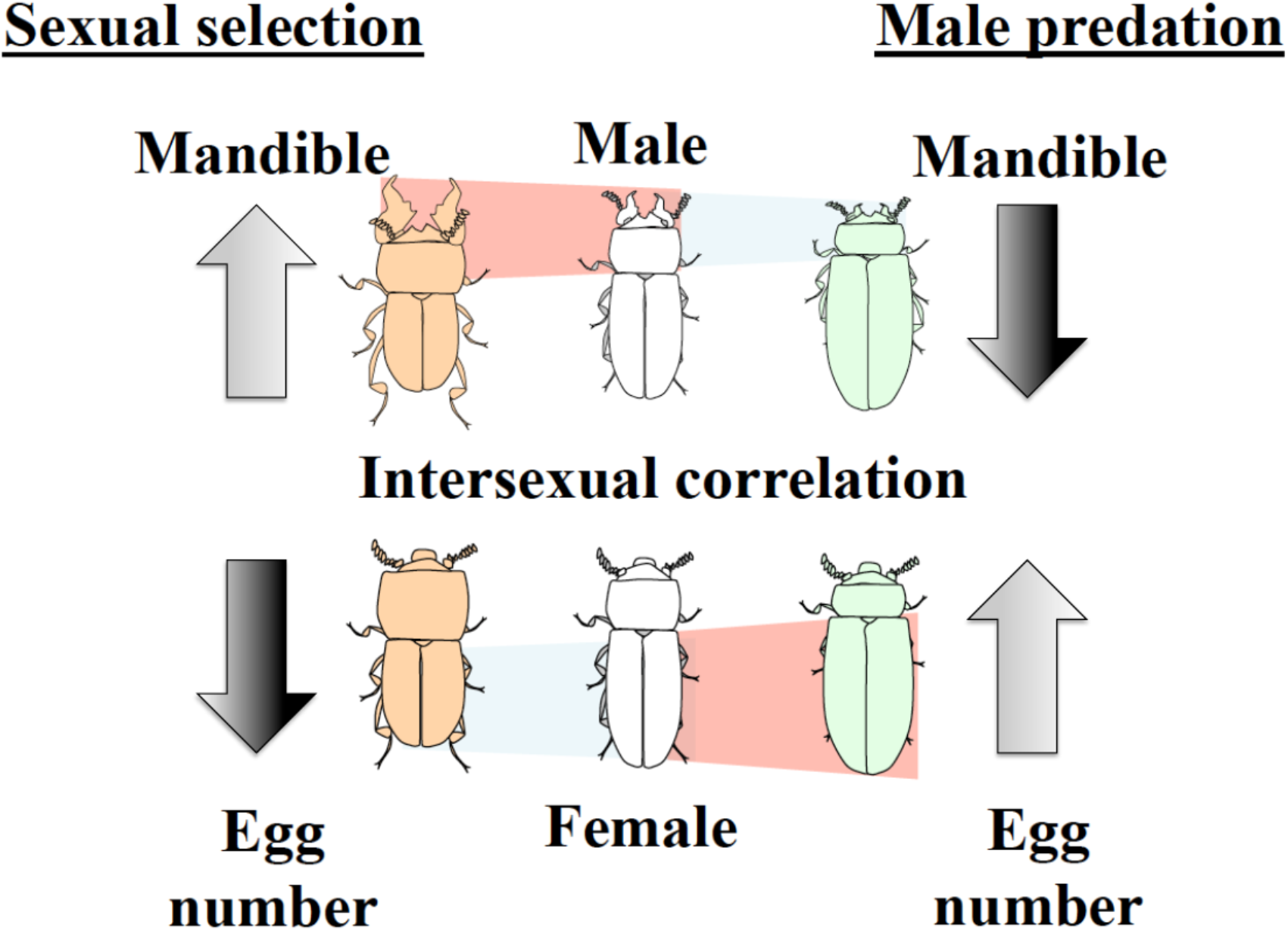
A diagrammatic representation of the male and female phenotypes resulting from male-limited predation in comparison to those resulting from sexual selection on males. Sexual selection (left images) results in enlarged male mandibles, which require a masculinized head and prothorax to operate effectively. This fore-body masculinization results in a smaller male abdomen and because of intersexual correlations for abdomen size, a smaller female abdomen and capacity for fewer eggs, even though females never develop mandibles. Male-limited predation selects against the masculinized phenotype, ultimately resulting in larger male and female abdomens, and hence more eggs and higher fitness females (images on the right).

## Discussion

Predation is frequently invoked as an evolutionary brake on the exaggeration of sexually selected traits and there are many studies consistent with this logic, usually documenting selection against larger characters or assessing macro-evolutionary patterns consistent with it (e.g. Zuk et al. 1993; Millar et al. 2006). Here we employed experimental evolution and directly demonstrate that male-specific predation not only reversed the exaggeration of a sexually selected trait, but additionally, this reversal resulted in higher female fitness. This boost to female fitness occurred because predators selecting against mandible enlargement in males results in a less masculinized phenotype (a larger abdomen), and because of shared genetic architecture across the sexes, this allows females to become more feminized and produce more offspring. We discuss these findings in turn.

When males were exposed to predation, they evolved smaller mandibles. Thus increased natural selection via predation reduced the size of a sexually selected trait, which is broadly consistent with theory (e.g. Lande 1981; Kirkpatrick 1982; reviewed in Andersson 1994). Previous work with *G. cornutus* suggests why this occurred. Mandible size is negatively phenotypically and genetically associated with locomotor activity (Fuchikawa and Okada 2013), and locomotor activity (running) is a predator escape mechanism in flour beetles (Miyatake et al. 2008). Reduced running and lower escape rates for males with large mandibles would explain the microevolutionary pattern we detected. Interestingly, there is no intersexual correlation for locomotion in this beetle (Fuchikawa and Okada 2013), so there was no expectation that predation on females should impact male running speed and hence morphology. Be that as it may, we clearly showed that natural selection reversed the evolutionary exaggeration of a sexually selected male trait.

Females from the male-predation populations evolved higher fitness (LRS) even though females were not directly exposed to predators. Harano et al. (2010) demonstrated that directly artificially selecting for larger (male) mandibles reduces female fitness. This occurred because selection for increased mandible size resulted in a more masculinized phenotype and this masculinization ripples through inferred intersexually-shared genetic architecture to increase the masculinization of the female phenotype, reducing female fitness. The intersexual genetic associations we documented here, especially the negative mandible-female LRS correlation, flesh out this explanation and confirm previous inference (Harano et al. 2010). Negative intersexual fitness associations are common (Arnqvist & Rowe 2005; Bonduriansky & Chenoweth 2009) because alleles conferring high fitness to one sex frequently lower fitness in the other (e.g. Chippindale et al. 2001; Rostant et al. 2015). In addition to any sexually antagonistic selection, male beetles with larger mandibles are also more aggressive toward females (Kiyose et al.

2015). Thus exposure to males with larger mandibles potentially reduces female LRS due to (misdirected) male attacks (Kiyose et al. 2015). Therefore there could be two avenues to increased female fitness when predators select against large male mandibles, a reduction in ontogenetic conflict load (intralocus sexual conflict load), or reduced aggression. In any case, predation caused an evolutionary reduction in mandible size and this resulted in increased female fitness. Thus male-biased predation indirectly selects for increased female quality.

The net population level effects of sexual selection and sexual conflict over optimal phenotypes are not clear (e.g. Kokko and Brooks 2003; Arnqvist & Rowe 2005), with for example, evidence that sexual selection can both increase and decreases population extinction rates (e.g. Doherty et al. 2003; Jarzebowska & Radwan 2010; Lumley et al. 2015). Additionally, intralocus sexual conflict (as documented in the flour beetle: Harano et al. 2010) is thought to constrain population adaptation because sexually antagonistic selection keeps each sex from its fitness optima (Rice 1992; Arnqvist & Rowe 2005). Our results suggest that predator purging of males with the largest sexual traits reduces intralocus (ontogenetic) sexual conflict costs, enhancing female reproductive performance, which should (all else being equal) increase population productivity. Relaxing other sexual conflict can also increase population fitness (e.g. Holland & Rice 1999; Martin & Hosken 2004). It is interesting to note that predators are usually seen as suppressing prey populations (Nelson et al. 2004), which can have indirect ecological benefits for prey competitors (e.g. Paine 1966). As we have shown, sex-biased predation can have analogous indirect effects intra-specifically. However, the indirect impacts we documented were unidirectional since female-biased predation did not alter female fitness or male sexual-trait size.

Negative intersexual correlations for fitness are an indicator of intralocus sexual conflict (Bonduriansky & Chenoweth 2009) - although this can be complicated by *Wolbachia* infection (Duffy et al. 2019) - and we find negative correlations for fitness surrogates here – mandible size determines male fitness and LRS reflects female fitness (e.g. Harano et al. 2010; Katsuki et al. 2012b; Okada et al. 2014). Regardless of arguments about fitness and fitness correlates, this negative association provides the genetic link to correlated responses to selection on mandible size previously documented (e.g. Harano et al. 2010) and to those we see here.

Overall this study provides direct evidence that predator-mediated natural selection can evolutionarily reverse the exaggeration of a sexually selected trait. This finding is consistent with a vast body of fundamental theory (e.g. Lande 1981; Kirkpartick 1982; Hall et al. 2000) and empirical evidence (e.g. Zuk et al. 2006; reviewed in Andersson 1994). We also reveal novel outcomes when natural selection targets sex-limited sexually selected characters, since predator removal of a male imposed conflict load increased female fitness. Thus sex-biased predation within a species can essentially mimic indirect ecological competition effects. Investigating the precise mechanistic detail of some of these findings is now required.

## Methods

### *Gnatocerus* stock culture

The *G. cornutus* beetle culture originated from adults collected in Miyazaki City (31° 54’N, 131° 25’ E), Japan, and has been maintained in the laboratory of the National Food Research Institute, Japan, for about 50 years on whole meal enriched with yeast as food. The stock is made up of 1500-2000 beetles per generation and maintained in plastic cups (diameter 95 mm, height 50 mm) with a standing density of between 300 and 400 beetles per cup (for a more detailed description of the stock culture, see Okada & Miyatake (2010)). This beetle is a stored product pest, and thus the laboratory conditions very closely mimic what have become their natural conditions. Virgin males and females were removed from the stock population as final instar larvae. Each larva was placed in one well of a 24-well tissue culture plate (Cellstar; Greiner Bio-One, Frickenhausen, Germany) until adult eclosion because pupation in *G. cornutus* is inhibited under high larval density (Okada & Miyatake 2010). After eclosion, both sexes were allowed to sexually mature for a period of 14 days prior to their use. We performed all rearing and experiments in a chamber maintained at 25°C, 60% relative humidity, and with a photoperiod cycle of 14:10 h light/dark. All experiments in this study follow this protocol unless stated otherwise.

### The predator

The assassin bug *Amphibolus venator* is predator of stored-product insect pests and preys on various stored-product insect pests including flour beetles (Pingale 1954; Nishi and Takahashi, 2002; Imamura et al. 2008). These predators are frequently found in stored product facilities, which are the habitat of *G. cornutus* (Nishi & Takahashi, 2002). The *A. venator* culture originated from adults collected in a storehouse in Urasoe City, Okinawa, Japan, and has been maintained in the laboratory for about 5 years. The stock was initiated and maintained at 200 bugs per generation and housed in plastic containers (230 mm × 150 mm × 80 mm) with a standing density of between 30 and 40 bugs per cup. Each nymph was given an excess of food (7 final instar larvae of *G. cornutus* per week). Each adult female was allowed to mate with a male chosen randomly and to lay eggs in order to maintain the predator stocks.

### *Gnatocerus* breeding design and estimation of quantitative genetic parameters

Using a full sib/half sib experimental design, males (sires) (N = 35) were randomly assigned to three virgin females (dams) (all collected from the stock population). Pairs were housed in a plastic container (17 mm diameter, 20 mm high) containing filter paper (17 mm diameter), and successful copulation was indicated by a stable end-to-end connection between the male and female. After mating, dams were immediately removed and individually placed in a plastic cup (70 mm diameter, 25 mm high) containing excess food (20 g). Each female was housed thus for two months to obtain offspring. All offspring from each female were reared to final instar (approximately 8 weeks). Three sons and three daughters per dam per sire were haphazardly chosen for measurement of male traits (mandible, body and abdomen size) and female traits (LRS, body and abdomen size) at 14 days after eclosion (N = 315 per sex) (trait measurement protocols below).

Data from the breeding design were then analysed using pedigree-based animal models fit in ASReml-R (Butler et al. 2017). First, to confirm the presence of additive genetic variance in each trait we fit a series of univariate animal models to (male) mandible, body and abdomen size, and to female (LRS). For each trait we compared the model fit to a reduced model with no additive genetic effects using a likelihood ratio test (LRT; adjusted for boundary conditions following Stram & Lee (1994)). We elected to combine male and female records for both body size and abdomen sizes as additional modelling provided little support for genotype-by-sex interactions (see results). However, for these traits a fixed effect of sex was included, as well as the (random) additive genetic effect since exploratory analysis showed sexual dimorphism in both traits (body size, males are 0.020 mg (SE 0.005) larger on average, t=3.58, P<0.001; abdomen size, males are 0.019 mg (SE 0.008) smaller on average, t=2.463, P=0.014)).

We then fitted a multivariate animal model to estimate genetic correlation (r_G_) structure among the four traits with fixed effects of sex on body size, abdomen size, as well as their heritability (h^2^). The residual covariance structure was modelled as an unstructured matrix (but note residual covariance between the sex limited traits of male mandible size and female LRS is not estimable from the data so was fixed to zero). We also ran reduced multivariate models with i) no genetic effects at all, and ii) a diagonal genetic variance matrix (i.e. genetic variance modelled on all traits but all genetic correlations assumed to equal zero) for comparison to the full model by LRT. This allows statistical inference at the level of the multivariate phenotype. We used estimated standard errors (SE) as a guide to nominal significance of pairwise genetic correlations (assuming approximate 95% CI are given by r_G_ ± 1.96SE).

### *Gnatocerus* experimental evolution protocol – sex specific predation

We first collected 900 male and 900 female *G. cornutus* from the stock culture and haphazardly generated 9 groups of 100 males and 100 females to establish three male-predation populations, three female-predation populations and three control (no predation) populations (generation 0). To simulate predation, 100 males (or females) were housed in a plastic container (150 mm diameter, 50 mm high) containing an excess of beetle food (45 g). Then, five adult female *A. venator* (20-35 day olds) were randomly collected from the predator culture and placed into the container and the males (females) were exposed to them for two weeks. We then selected 10 of the males (females) that survived the two weeks to act as sires (dams) of the predation treatments - 10 opposite sex individual were also taken/population to act as the non-selected dams (sires) that were not exposed to predation. We note that survival rate during this predation protocol was approximately 20%. To propagate control populations, 10 males and 10 females were haphazardly selected per population to act as sires and dams. For each population/treatment the 10 males and females were placed in a plastic cup (diameter 95 mm, height 50 mm) with 70 g of medium for two months, with males able to mate with females and females were allowed to lay eggs, until final instar larvae were obtained. Final instar larvae were collected to obtain the adults for subsequent generations. When the adults reached 14 days old, 100 males and 100 females per population were randomly selected to (potentially) seed the next generation, and in the predation treatments, exposed to predators as above. We then selected surviving animals as above and repeated for 8 generations. Additionally, we also collected 50 males and 50 females per population from generation 1 to 7 to assess mandible, abdomen and body size responses to selection. At generation 8, 20 males and 20 females per population were haphazardly collected for measurement of male traits (mandible, body and abdomen size) and female traits (LRS, body and abdomen size) (N = 180 per sex) (trait measurement protocols below).

### Male trait measurement

We measured overall body mass and the posterior body mass (i.e., mesothorax, metathorax, and abdomen) as an abdomen size indicator (see Harano et al. 2010). Briefly, each male was frozen at −20 °C immediately after adult emergence. Mass measures were obtained to the nearest 0.01 mg on an electronic balance (Mettler-Toledo AG, Laboratory and Weighing Technologies). The mandible length (±0.01 mm) of each male was measured (±0.01 mm) using a dissecting microscope monitoring system (VM-60; Olympus, Tokyo, Japan). Each specimen was positioned so that its longitudinal and dorsoventral axes were perpendicular to the visual axes of the microscope eyepiece (see Okada & Miyatake, 2010 for landmarks).

### Female trait measurement

To obtain LRS (lifetime reproductive success: our fitness proxy) each female (14 days post-eclosure) was individually paired with a haphazardly selected male from the stock culture. After copulation, each female was maintained in a plastic cup (70 mm diameter, 25 mm high) containing an excess of with food (20 g) for two months and allowed to lay eggs. This schedule was chosen because most eggs are laid by females within two months of mating (Tsuda and Yoshida 1984), and thus this is an accurate index of LRS (Katsuki et al. 2012ab). To measure the LRS of each female, we counted all adults that emerged in the third month after pairing. After the laying period, each female was frozen at −20 °C. Subsequently, the whole and posterior of the body (body size and abdomen size) were weighed with the electronic balance (as above).

Apart from the genetic parameter estimation with an animal model (using ASReml-R as described above), all analyses were conducted using JMP for Windows version 8 (SAS Institute 2008). We used population as the unit of replication (= 9 DF max.) with single fixed factor (with 3 levels: the experimental treatments) GLMs for each trait to test for effects of experimental evolution, with post-hoc testing for factor-level differences (note we did not have sufficient DF for a multivariate analysis). Results are as reported even after (conservative: Nakagawa 2004) sequential Bonferroni correction.

## Acknowledgements

We thank Stu Bearhop, Sasha Dall and Dave Hodgson for discussion. DJH was supported by the Leverhulme Trust.

## Supplementary Information

### Estimated fixed effects and heritabilities from univariate animal models of each trait

Body mass and abdomen mass were treated as single traits rather than sex-specific ones, with sex included as a fixed factor. Statistical inference on fixed effects is by conditional F test while the presence of additive genetic variance was tested by LRT comparison to a reduced model assuming twice the difference in model log-likelihoods is distributed as a 50:50 mix of X^2^_0_ and X^2^_1_.

|  | Fixed effects |  |  |  |  | Random effects |  |  |
| --- | --- | --- | --- | --- | --- | --- | --- | --- |
| Trait | Effect | Estimate (SE) | DF | F | P | $h^2$ (SE) | $X^2_{0,1}$ | P |
| Body mass | mean | 2.701 (0.005) | 1,50.9 | 318800 | <0.001 | 0.343 (0.085) | 33.95 | <0.001 |
|  | Sex (male) | 0.020 (0.005) | 1,528.7 | 15 | <0.001 |  |  |  |
| Abdomen mass | mean | 1.843 (0.008) | 1,57.2 | 55940 | <0.001 | 0.508 (0.096) | 58.11 | <0.001 |
|  | Sex (male) | -0.019 (0.007) | 1,523.0 | 8 | 0.005 |  |  |  |
| Male mandible length | mean | 0.396 (0.004) | 1,44.0 | 10050 | <0.001 | 0.286 (0.121) | 8.50 | 0.002 |
| Female LRS | mean | 141.873 (1.714) | 1,41.8 | 6849 | <0.001 | 0.165 (0.108) | 3.234 | 0.036 |

### Tests for genotype-by-sex interaction in body and abdomen mass

Shown are likelihood ratio test (LRT) comparisons of a simple univariate animal model (including a fixed effect of sex) to once in which genotype-by sex interaction is modelled. Twice the difference in log-likelihoods is assumed to be distributed as X^2^ with 2DF. Also shown are estimates of genetic variance (V_A_) under the simple model and the sex-specific genetic variances (V_Af_, V_Am_) and the cross-sex genetic correlation (r_Gmf_) under the expanded model allowing genotype-by-sex interaction. Standard errors are shown in parentheses where available. Note that to keep the genetic variance-covariance matrix in allowable parameter space (i.e. positive definite) rGmf was bound to (effectively) +1 in both expanded models and no SE is estimated as a consequence. For body mass the improvement to model fit is marginally non-significant when the genotype-by-environment interaction is included. To the extent this might reflect real differences in sex-specific genetic architecture the pattern is driven by apparent differences in VA across the sexes (rather than deviation from r_Gmf_=1). Consequently, we assuming an absence of genotype-by-sex interaction for this trait, we also note that the estimated heritability presented in the main manuscript remains valid (as an average across the sexes) even if this assumption is incorrect.

|  | Model comparison |  | Simple animal model | Expanded model with genotype-by-sex interaction |  |  |
| --- | --- | --- | --- | --- | --- | --- |
| Trait | $X^2_2$ | P | $V_A$ (SE) | $V_{Af}$ (SE) | $V_{Am}$ (SE) | $r_{\text{Gmf}}$ |
| Body mass | 5.49 | 0.075 | 0.0017<br>(0.0005) | 0.0026<br>(0.0006) | 0.0013<br>(0.0005) | 0.9999 |
| Abdomen mass | 0.20 | 0.915 | 0.0050<br>(0.0012) | 0.0053<br>(0.0014) | 0.0047<br>(0.0013) | 0.9999 |

